# Sex-biased gene expression at single-cell resolution: Cause and consequence of sexual dimorphism

**DOI:** 10.1101/2022.11.08.515642

**Authors:** Iulia Darolti, Judith E. Mank

## Abstract

Gene expression differences between males and females are thought to be key for the evolution of sexual dimorphism, and sex-biased genes are often used to study the molecular footprint of sex-specific selection. However, gene expression is often measured from complex aggregations of diverse cell types, making it difficult to distinguish between sex differences in expression that are due to regulatory rewiring within similar cell types and those that are simply a consequence of developmental differences in cell type abundance. To determine the role of regulatory versus developmental differences underlying sex-biased gene expression, we use single-cell transcriptomic data from multiple somatic and reproductive tissues of male and female guppies, a species which exhibits extensive phenotypic sexual dimorphism. Our analysis of gene expression at single-cell resolution demonstrates that non-isometric scaling between the cell populations within each tissue and heterogeneity in cell type abundance between the sexes can influence inferred patterns of sex-biased gene expression by increasing both the false-positive and false-negative rates. Moreover, we show that at the bulk level, the subset of sex-biased genes that are the product of sex differences in cell type abundance can significantly confound patterns of coding-sequence evolution. Taken together, our results offer a unique insight into the evolution of sex-biased gene expression and highlight the power of single-cell RNA-sequencing in disentangling between genes that are a cause as opposed to a consequence of sexual dimorphism.

## Introduction

Males and females of the same species often exhibit striking differences in a broad range of phenotypic traits, despite sharing the majority of their genome. It is therefore widely assumed that transcriptional changes between the sexes are key to the evolution of sexual dimorphism (Grath and Parsch 2016; Mank 2017). Indeed, previous work has shown that the magnitude of transcriptional dimorphism scales with the level of phenotypic dimorphism across a large proportion of the genome (Pointer et al. 2013). Consistent with their sex-specific roles, reproductive organs are by far the most transcriptionally dimorphic tissues, however there is substantial variation in sex differences in expression across somatic tissues as well (Yang et al. 2006; Mank et al. 2007; Ma et al. 2018; Khodursky et al. 2020; Oliva et al. 2020). Ontogenetic studies have also revealed that the degree of sex-biased gene expression amplifies over the course of development, reflecting the increase in phenotypic dimorphism (Mank et al. 2010; Magnusson et al. 2011; Perry et al. 2014; Ingleby et al. 2015; Djordjevic et al. 2022).

Although the above-mentioned studies provide evidence for a correlation between transcriptional and phenotypic dimorphism across multiple levels of biological diversity, few functional validation experiments have tested the causal effect of sex-biased gene expression (Abzhanov et al. 2006; Khila et al. 2012; Chen et al. 2015; Galouzis and Prud’homme 2021; Toubiana et al. 2021). These functional assays are often limited by the sheer number of sex-biased genes, the polygenic nature of traits, or simply because the necessary genetic tools are lacking, a particular problem for non-model organisms, which often present the most striking phenotypic differences. As such, it remains unclear to what extent are sex differences in expression a cause, as opposed to a consequence, of sexual dimorphism.

Many studies have also used sex-biased gene expression as a way to measure the footprint of sex-specific selection within the genome, however, this approach has produced some discordant results (Grath and Parsch, 2016; Mank 2017). Male-biased genes in many species tend to exhibit higher rates of evolution at both the coding sequence and the expression level (Ranz et al. 2003; Khaitovich et al. 2005; Sharma et al. 2014; Lipinska et al. 2015; Whittle and Extavour 2019; Lichilin et al. 2021). While early work in *Drosophila melanogaster* has interpreted this as the result of stronger sexual selection acting in males (Proschel et al. 2006; Sawyer et al. 2007), studies in other species have found that such accelerated patterns of evolution are instead more consistent with relaxed constraint (Gershoni and Pietrokovski et al. 2014; Harrison et al. 2015; Sayadi et al. 2019; Dapper and Wade, 2020; Djordjevic et al. 2021), and experimental evolution results were somewhat mixed (Hollis et al. 2014; Veltsos et al. 2017; Li Richter and Hollis 2021).

Some of this discordance in results may be due to the way we measure expression. Most sex-biased gene expression studies have so far relied on bulk RNA-sequencing, comparing expression between organs (Harrison et al. 2015; Whittle and Extavour 2019), whole body parts comprising various tissues such as heads or abdomens (Standage et al. 2016; Immonen et al. 2017), or even entire organisms (Ranz et al. 2003; Hollis et al. 2014; Stuglik et al. 2014; Djordjevic et al. 2022), These samples represent complex aggregates of diverse cell types that may have variable expression profiles. In many species, males and females exhibit dimorphism in the relative size of their constituent body parts (Badyaev 2002), and allometric scaling could influence perception of gene expression differences in studies where whole organisms are used for RNA preparation (Montgomery and Mank 2016). In a similar way, whole tissue expression studies may be affected by the heterogeneity in cell type abundance or composition (Fair et al. 2020; Hunnicutt et al. 2021; Fuess and Bolnick 2021; Price et al. 2022), and there is indeed evidence that sex differences in cell type populations exist for many tissues (Mank and Rideout 2021). For example, differential rates of cell proliferation between males and females seem to underly the development of several sexually dimorphic ornamental traits, such as caudal fins in *Xiphophorus* (Schartl et al. 2021; Powell et al. 2021) and horns in rhinoceros beetles (Emlen et al. 2012).

Single-cell transcriptomics (scRNA-seq) offers the possibility to avoid the challenges posed by measuring expression from a heterogeneous tissue by instead comparing expression level between samples across equivalent cell populations. Here we leverage these recent advances in scRNA-sequencing to determine to what extent is sex-biased gene expression the result of sex differences in cell type abundance as opposed to regulatory differences between similar cells, and to assess how this impacts inferences of evolutionary divergence.

## Methods

### Tissue collection and dissociation

We sampled reproductively mature male and female guppies from our laboratory population. All fish were raised at a water temperature of 26°C with a 12:12 light:dark schedule, and fed a daily diet of flake food and live Artemia brine shrimp. Fish were euthanized with a pH-neutralized MS222 solution and dissected immediately. We dissected liver, heart, tail skin (removing any muscle and scales) and gonad (testis in the case of males, and ovaries excluding mature oocytes in the case of females) tissues and immediately placed them in PBS solution (Corning) on ice. We obtained three replicates for each tissue type and sex, and every replicate contained a non-overlapping pool of five tissues in order to ensure sufficient material for single-cell dissociation, resulting in a total of 24 samples.

Tissues were incubated at 30°C in a solution of 4mg/ml collagenase type I (Sigma-Aldritch) and 4mM CaCl_2_, gently pipetting every 3 minutes using a wide-bore pipette tip, until digested. Dissociated tissues were then filter-strained using a Flowmi 40μm cell strainer. We centrifuged the samples at 300 rcf for 3 minutes, removing the supernatant and resuspending the cell pellet in PBS containing 0.04% bovine serum albumin (BSA) (Corning). Samples were centrifuged and resuspended in PBS + 0.04% BSA twice and immediately placed on ice before further processing.

### Single-cell library preparation and sequencing

We mixed 10μl of cell solution with 10μl of the exclusion dye trypan blue 0.4% (Invitrogen) and estimated cell viability and concentration using a Countess II automated cell counter (ThermoFisher). An estimated 8,000 cells from each sample were then loaded onto individual lanes of a 10X Genomic Chromium Controller and barcoded 3’ single cell libraries were prepared using the 10X Genomics Chromium Next GEM Single Cell 3’ kit v3.1 following the manufacturer’s instructions (#CG0000204). We assessed the quality and concentration of cDNA and libraries using an Agilent 4200 TapeStation and the Agilent High Sensitivity D5000 ScreenTape system. Libraries were sequenced on an Illumina NovaSeq 6000 sequencer, with an average sequencing depth of 20,000 read pairs per cell.

### scRNA-seq data processing

We used CellRanger v5.0.1 with the ‘mkref’ function (Zheng et al. 2017) to build a reference index using the Ensembl *P. reticulata* genome (GCA_000633615.2) and annotation (release-103) files. Using the CellRanger v5.0.1 ‘count’ function, we then aligned sequencing reads from fastq files to the reference index, identified cell-associated barcodes and extracted gene-by-cell count matrices. Data filtering and downstream analyses were performed using Seurat v4.1.0 (Satija et al. 2015) in R v4.0.5 (R Core Team 2021). We filtered the count data by keeping genes expressed in at least three cells and cells with at least 100 expressed genes. Raw counts were then normalized and scaled for each sample to account for differences in sequencing depth per cell using the ‘SCTransfrom’ function (Hafemeister and Satija 2019). We used the function ‘doubletFinder_v3’ from the DoubletFinder v2.0.3 package (McGinnis et al. 2019) in R to identify and remove doublets, which are the result of random encapsulation of more than one cell within the single-cell bead as part of the microfluidics process.

### scRNA-seq data clustering

We performed the principal component analysis (PCA) using the ‘RunPCA’ function and identified the significant PCs using the ‘ElbowPlot’ function. We then constructed the nearest-neighbor graph with the ‘FindNeighbors’ function and performed the graph-based clustering with the ‘FindClusters’ function. We used the clustree v0.4.4 package in R to guide the decision of the optimal resolution to choose for the clustering analysis (Zappia and Oshlack 2018). Clusters were then visualized by using uniform manifold approximation and projection (UMAP) embedding with ‘RunUMAP’. For each tissue, we used the ‘FindAllMarkers’ function in Seurat to identify differentially expressed genes for each cell cluster. We then annotated the major cell types within each cluster based on marker gene information from the Zebrafish Cell Landscape (Jiang et al. 2021; Wang et al. 2022) and other published work (Morrison et al. 2021; Liu et al. 2022; Qian et al. 2022).

### Differential cell type abundance analysis

For each tissue type, we tested for significant differences between male and female samples in the abundance of the identified cell types. We first quantified the number of cells assigned to each cell type and sample. We then merged counts for each sex and calculated the proportion of cells of each cell type out of the total number of cells. We added 1e-10 to each proportion value to avoid infinitely high numbers associated with log_2_ 0, and calculated the female-to-male fold change (FC) in cell type abundance as log_2_ (female proportion/male proportion). To identify significant differences in cell type abundance between the sexes we computed two-proportions z-tests (*p* < 0.01) with the ‘prop.test’ function in R.

### Sex-biased gene expression analysis

To identify sex-biased genes at the cell-level, for each identified cell type, we aggregated raw counts across all cells to the sample level using the ‘aggregate.Matrix’ function in R. We then used DESeq2 to normalize the count data accounting for differences in library size between samples and applying a log2-transformation with the ‘rlog’ function (Love et al. 2014). Lastly, we performed the differential expression analysis using the ‘DESeq’ function. Sex-biased genes were called based on |log_2_ FC| >= 1 and a false discovery rate (FDR) adjusted *p* value < 0.05 to correct for multiple testing. To identify sex-biased genes at the bulk-level we followed the same steps described above, but instead aggregated counts across all cells from all clusters together to obtain a single expression value for each gene and sample.

### Rates of coding-sequence evolution analysis

We obtained coding sequences from the outgroup species *Gambusia affinis* (ASM309773v1), *Xiphophorus maculatus* (Xipmac4.4.2) and *Oryzias latipes* (MEDAKA1) from Ensembl 104 and extracted the longest isoform for each gene. We used reciprocal BLASTn v2.7.1 (Altschul et al. 1990) with an e-value cut-off of 10e-10 and a minimum percentage identity of 30% to determine orthology across the *P. reticulata* genes and outgroup sequences. For genes with multiple blast hits, we chose the top hit based on the highest BLAST score.

We used *O. latipes* (MEDAKA1) protein-coding sequences from Ensembl 104 and BLASTx v2.3.0 with an e-value cut-off of 10e-10 and a minimum percentage identity of 30% to obtain open reading frames. We excluded orthogroups without BLASTx hits or valid protein-coding sequences. We aligned orthologous gene sequences with PRANK v170427 (Löytynoja and Goldman 2008), and filtered alignments to remove gaps. We also masked poorly aligned and error-rich regions with SWAMP (Harrison et al. 2014) with a threshold of 4 misalignments in a window size of 5bp and a minimum sequence length of 100bp.

To obtain divergence estimates, we used the branch model (model=2, nssites=0) in the CODEML package in PAML v4.8 (Yang 2007). Genes with *d*_S_ > 2 were removed from subsequent analyses to avoid inaccurate divergence estimates due to mutation saturation and double hits (Axelsson et al. 2008). We divided genes into different sex-bias categories (see Fig. 3) and extracted the number of nonsynonymous (*D*_N_) and synonymous substitutions (*D*_S_) and the number of nonsynonymous (N) and synonymous (S) sites. For each group of genes, we then calculated the mean rate of nonsynonymous substitutions (*d*_N_) and mean rate of synonymous substitutions (*d*_S_) as the number of substitutions across all genes divided by the number of sites (*d*_N_=*D*_N_/N; *d*_S_=*D*_S_/S). We used bootstrapping with 1,000 replicates to determine the 95% confidence intervals for divergence estimates in each gene group, and tested for differences in *d*_N_, *d*_S_ and *d*_N_/*d*_S_ estimates between the different gene groups based on 1,000 replicates permutation tests.

## Results

We generated 24 scRNA-seq datasets from skin, heart, liver and gonad tissue from adult male and female guppies, with three replicates for each sex and tissue. Guppies are highly sexually dimorphic, displaying sex differences in size (Houde 1997), sexual ornaments (Reznick and Endler, 1982), life history (Reznick and Endler, 1982), and behavior (Reznick 1989), among other traits, and we chose these four tissues to reflect a range of phenotypic dimorphism. Following quality control and filtering (see Methods), we recovered between 503-3,747, 1,163-8,966, 4,985-6,835 and 1,492-7,973 cells, and 9,297, 9,860, 11,478, and 13,617 genes expressed in skin, heart, liver and gonad tissues, respectively. Following UMAP dimensionality reduction and marker-based annotation we identified 13, 11, 9 and 8 distinctly expressed cell clusters for skin, heart, liver and gonad, respectively (Fig. S1; Fig. S2). These clusters are representative of most major cell types found in these tissues (Jiang et al. 2021; Wang et al. 2022).

### Sex differences in cell type abundance across tissues

We examined each tissue for differences in cell composition between the sexes based on the estimated abundances of each of the identified cell types. We found significant sex differences (*p* < 0.01, two-proportions z-tests) in abundance for many cell populations (Fig. 1; Table S1), the most extreme differences being present in the reproductive tissue, where most of the cell types are sex-limited (Fig. 1D, Fig. S3). Notably, the vast majority of cells in the male gonad are related to sperm, while the female gonad represents a broader mix of gametic and somatic cells (Fig. S3). However, we observe a range of dimorphism in cell composition among somatic tissues as well. The liver plays a critical role in several physiological processes, including hormonal regulation, metabolism, digestion and immune response (Bruslé and Anadon, 1996), and shows marked sex differences (Marcos et al. 2015; Ullah et al. 2021). Consistent with the more sexually dimorphic nature of this somatic tissue, we found extensive sex differences in cellular composition, with all but one of the identified cell types showing a significant sex-bias in abundance (Fig. 1C). Although the heart and skin tissues are less sexually dimorphic compared to the liver, we also recovered sex differences in abundance for more than half of the cell populations in these two tissues (Fig. 1A, B). Markedly, guppies exhibit striking sexual dimorphism in skin pigmentation, with male-specific ornamental color patterns (Reznick and Endler, 1982), which we also find reflected here through the strong male-biased abundance of melanocytes in males (Fig. 1).

**Figure 1.**
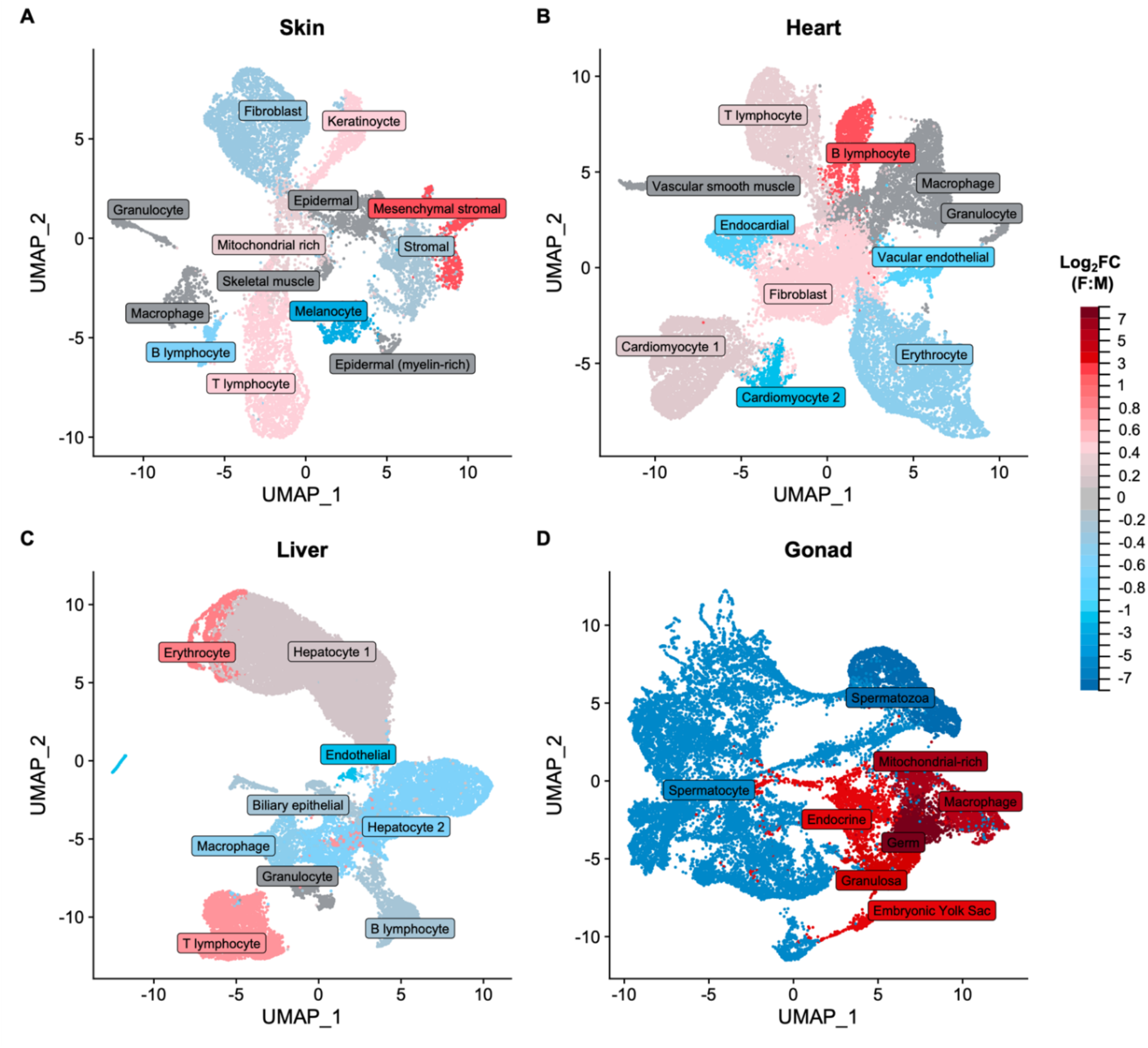
Differential cell type abundance between males and females in skin (A), heart (B), liver (C) and gonad (D) tissues. Cell types that are significantly more abundant in females are shown in red, those that are significantly male-biased in abundance are shown in blue, while unbiased cell types are in gray. Significance based on two-proportions z-tests (*p* < 0.01).

### Sex-biased gene expression at bulk- and cell-level

For each tissue, we aggregated expression measures across all cells to obtain a single value for each gene and sample, thus reflecting expression at the bulk tissue level. Based on this, we identified genes with a sex-biased expression profile (|log_2_ fold change| >= 1, FDR-corrected *p* value < 0.05). Consistent with previous estimates in guppies (Sharma et al. 2014), we find that at the bulk level male and female gonads exhibit strong patterns of sex-biased gene expression, with 9,524 genes identified as differentially expressed, representing 70% of all genes expressed in the gonads (Fig. 2D, Table S2). Top male-biased genes included those encoding for spermatogenesis processes (*march11, morn3, spata18, spatc1l*), ciliary- and flagellar-associated proteins (*cfap52, ropn1l, tekt1*), male-specific development (*dmrt1*) and calcium-binding proteins (*efcab2*), while female-biased genes encoded for, among others, zona pellucida glicoproteins (*zp2l2, zp3f*.*1, zp3d*.*2*), oocyte-specific proteins (*zar1, zar1l*) and ovarian folliculogenesis processes (*gdf9, cx43*.*4, pcdh18a*) (Table S2). Across the investigated somatic tissues, liver was the most transcriptionally dimorphic tissue, with 349 sex-biased genes, more than twice as many compared to the skin (114) and heart (155) tissues (Fig. 2A-C). The liver has been shown to be one of the more transcriptionally dimorphic somatic tissues in other fish species as well (Taboada et al. 2012; Zheng et al. 2013; Rose et al. 2015; Qiao et al. 2016). In fish and in other oviparous vertebrates, the liver carries an important role in the vitellogenesis process of egg yolk protein synthesis, transport and uptake in the maturing oocyte (Arukwe and Goksøyr 2003; Hara et al. 2016), and, in line with this, we also find vitellogenins (*vtg1, vtg2, vtg3*) to be some of the top female-biased genes in the liver (Table S2). While skin and heart tissues express fewer sex-biased genes, we also note highly male-biased genes in skin with a role in pigmentation (*fhl2a, pnp4a*).

**Figure 2.**
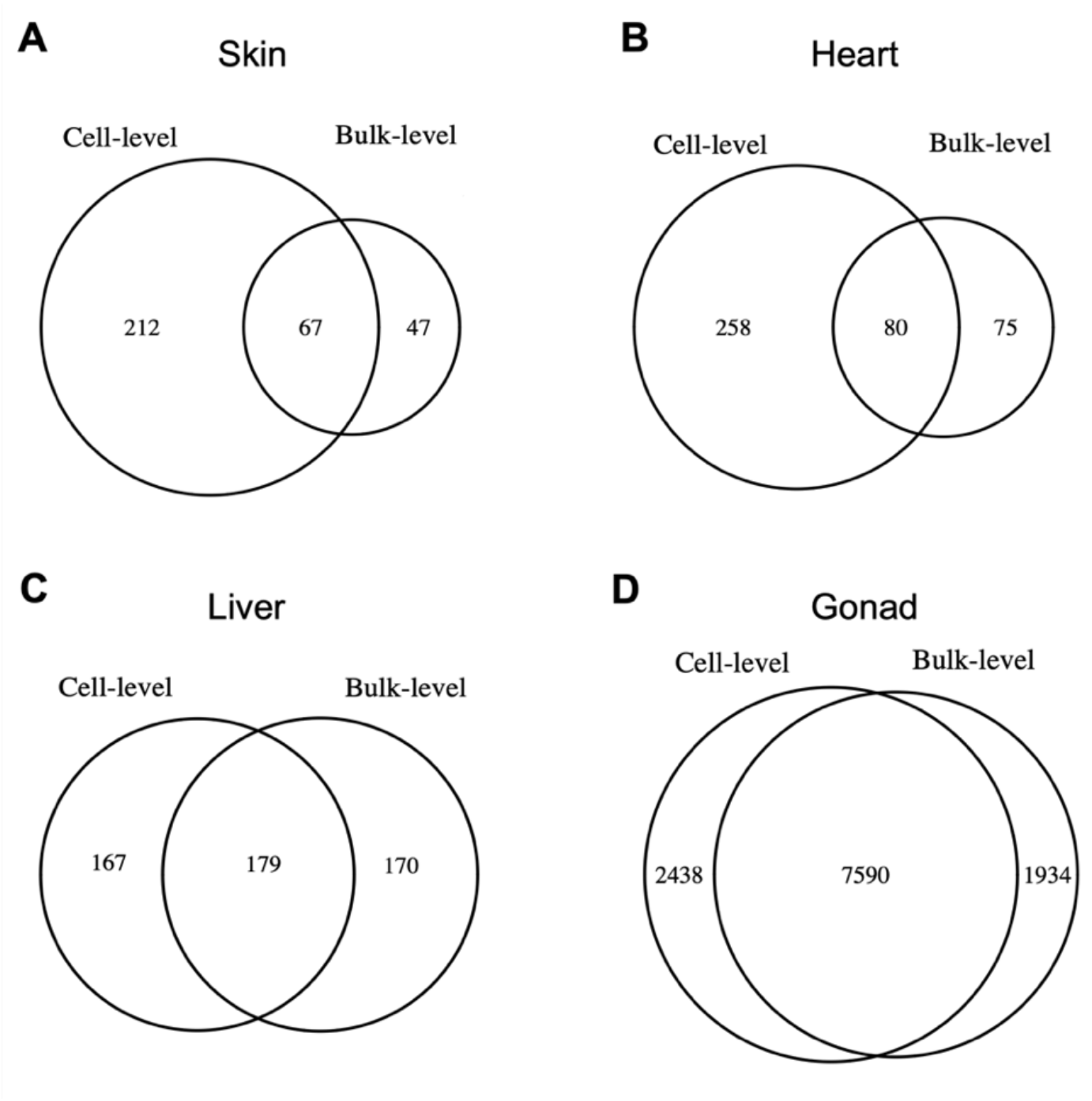
Number of genes showing sex-biased gene expression at the cell- and bulk-level in skin (A), heart (B), liver (C) and gonad (D) tissues. Numbers at the “Cell-level” represent the union of genes identified as sex-biased across all the different cell types. Significant differential expression between the sexes was based on |log2 FC| >= 1 and an FDR-corrected *p* value < 0.05.

We next compared patterns of differential gene expression between males and females at the whole tissue level with those at the cell level. As such, in addition to the aggregated bulk-level expression, for each identified cell type we aggregated expression data across all cells to identify sex-biased genes in each cell-type population. Overall, we discovered more sex-biased genes at the cell level than at the bulk level in all tissues, except for liver, however what is even more striking is that across the somatic tissues between 49-77% of the identified sex-biased genes at the cell level have unbiased expression profiles at the bulk level (Fig. 2). By contrast, in the gonad only about 24% of the differentially expressed genes at the cell level show no sex differences in expression at the bulk level. These results indicate that, outside of the reproductive tissues, differential gene expression analyses based on bulk RNA-sequencing data are limited in their ability to comprehensively identify sex-biased genes. Nonisometric-scaling relationships between cellular subcomponents of a tissue may cause patterns of differential expression in low abundance cell types to be concealed when expression is aggregated across various cell types in standard bulk RNA-sequencing experiments, thus creating a false-negative inference of sex-biased gene expression.

Moreover, in all somatic tissues, many genes are characterized as sex-biased at the bulk level but not at the cellular level. We hypothesized that these differential expression patterns are not due to regulatory rewiring within cell types but instead a consequence of differences in cell type abundance between males and females. Indeed, these genes tend to have a significantly lower magnitude of expression fold change (Fig. S4) and are more highly expressed in cell types that exhibit significant sex differences in abundance (Fig. S5). In skin, genes that are female-biased at the bulk level only are predominantly expressed in the mesenchymal stromal cell population which is more abundant in females. Similarly in liver, genes that are female- and male-biased in expression at the bulk level but not at the cell level have a significantly higher expression in the female-abundant T lymphocytes and the male-abundant endothelial cells, respectively. Although in the gonad a much smaller percentage of genes are sex-biased only at the bulk level, these too appear to be the result of sex differences in cell type abundance (Fig. S5). These results show that heterogeneity in cellular scaling relationships, due to developmental differences in cell proliferation, between males and females can influence perceived patterns of differential gene expression in the absence of regulatory changes.

### Rates of evolution of sex-biased genes

We tested whether the cause of sex-biased gene expression influences estimates of coding-sequence evolution. When considering all differentially expressed genes at the bulk level, we found that male-biased genes in the liver and gonad and female-biased genes in the skin, liver and gonad evolve significantly faster than unbiased genes (Fig. 3). These patterns are primarily driven by higher rates of nonsynonymous substitutions for the sex-biased genes in each group compared to the unbiased genes (Table S3). However, separating these sex-biased genes into those that are likely the result of sex differences in cell type abundance and those that are, at least in part, due to regulatory differences between males and females produced notable differences (Fig. 3). Specifically, the magnitude of divergence between unbiased and sex-biased genes was significantly more accentuated for genes that are sex-biased at both the bulk- and the cell-level. These loci are more likely to be underlying sexually dimorphic phenotypes as they also exhibit elevated log_2_FC estimates (Fig. S4). On the other hand, genes that are sex-biased at the bulk-level only, and thus the outcome of differential tissue composition between the sexes, showed similar rates of coding-sequence evolution to unbiased genes, further suggesting that they are not main targets of selection. For the gonad, this is particularly evident, as the vast majority of male-biased genes are related to sperm production (Fig. S3), while female-biased genes in the gonad represent a mix of somatic and gametic cell types. False-positive differentially expressed genes can therefore obscure true patterns of evolutionary divergence for sex-biased genes at the bulk level.

**Figure 3.**
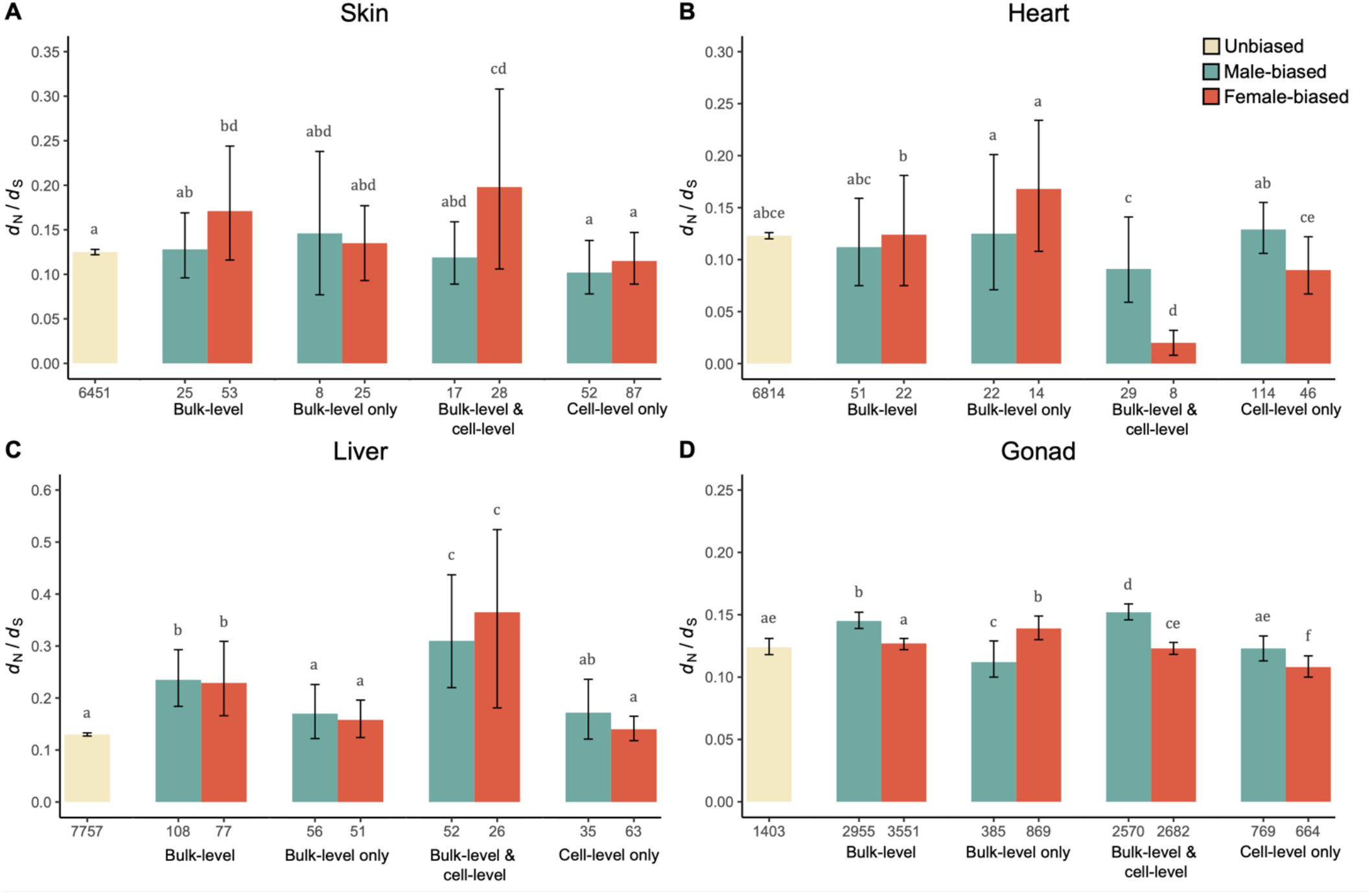
Rates of coding-sequence evolution (*d*_N_/*d*_S_) for genes in skin (A), heart (B), liver (C) and gonad (D). Shown are estimates of divergence for genes that are unbiased at both the bulk level and cell level (yellow), all genes identified as male-biased (green) and female-biased (orange) at the bulk level (Bulk-level), genes with a male-biased and female-biased expression at the bulk level but that are unbiased in every cell type within that tissue (Bulk-level only), genes that show a sex-biased expression profile both at the bulk level and in at least one of the identified cell types in that tissue (Bulk-level & cell-level), and sex-biased genes at the cell level but not at the bulk level (Cell-level only). Numbers on the x-axis represent the number of genes in each group. Different letters above bars indicate significant differences among groups (*p* < 0.05).

## Discussion

To determine the relative role of regulatory versus developmental differences underlying sex-biased gene expression, we use single-cell transcriptomic data from multiple somatic and reproductive tissues of male and female guppies, a species which exhibits extensive phenotypic sexual dimorphism. Sex-biased genes can be a cause of sexual dimorphism, as is often assumed in studies that use them to study the molecular genetic footprint of sexual selection and sexual conflict (e.g., Harrison et al. 2015; Mank 2017; Sayadi et al. 2019). Alternatively, sex-biased genes may be a consequence of developmental differences in cell proliferation between the sexes that result in differences in cell type abundances. Sex differences in cell populations are known to exist in many tissues (Mank and Rideout 2021), and differential rates of cell proliferation underly the development of several sexually dimorphic ornamental traits (Emlen et al. 2012; Schartl et al. 2021; Powell et al. 2021). The question of whether sex-biased genes are a cause or consequence of sexual dimorphism is critical, as the former might be subject to differences in sex-specific selection and therefore useful in studies of sexual selection, sexual dimorphism and sexual conflict, while the latter are largely a consequence of differences in developmental programming.

Our analysis of gene expression at single-cell resolution illustrates that cellular heterogeneity within tissues and allometric scaling differences of cell types between males and females can have a major influence on inferred patterns of sex-biased gene expression. Such scaling effects can generate both false-negative detection of differentially expressed genes, which may preponderantly concern genes that are expressed in low abundance cell types, as well as false-positive patterns of differential gene regulation, as is the case for genes that are sex-biased at the bulk tissue level but not at the cell level. These false-positive results have the potential to affect inferences on the evolution of sex-biased gene expression in several ways.

Sex differences in gene co-expression networks are thought to contribute to sexually dimorphic phenotypes and potentially alleviate sexually antagonistic selection (Sutherland et al. 2019; Lopes-Ramos et al. 2020; Rago et al. 2020). Yet fundamental differences in gene regulatory networks can exist between cell types and variation in cell type abundance between the sexes can affect gene co-expression measurements (Ribeiro et al. 2022). Allometry can also be a confounding factor in estimates of rates of coding-sequence evolution for sex-biased genes, as we show that false-positive sex-biased genes diminish the signal of elevated rates of divergence for both male- and female-biased genes. Variation in the fraction of genes that are erroneously classified as sex-biased due to sex differences in tissue composition may explain some of the discordance in patterns of rapid rates of coding-sequence evolution and sex-specific selection observed across studies (Ranz et al. 2003; Khaitovich et al. 2005; Mank et al. 2010; Whittle and Johannesson 2013; Harrison et al. 2015; Lipinska et al. 2015).

The degree of sex differences in cell type abundance likely varies across species as a function of phenotypic sexual dimorphism, and this can influence patterns of rapid turnover of sex-bias across species. We observed large differences across tissues in cell type abundance between the sexes (Table S1, Fig. 1, Fig. S3), with the liver showing the least overall fold change in cell type abundance, greater levels in the skin and heart, and the greatest observed in the gonad. In some cases, cell type abundance differences were consistent with visible phenotypic differences, such as the male-bias in melanocytes in male skin, as might be expected from male coloration. However, many cell types with sex differences might not necessarily be predicted from phenotypic differences, such as the male-bias in heart cardiomyocytes, or the female-bias in skin mesenchymal stromal cells.

Notably, spermatocytes and spermatozoa combined make up 97% of male gonad cells. In contrast, female gonad is comprised of only 20% germ cells, and the rest a mix of somatic cells. This means that bulk comparisons between male and female gonads or whole bodies, where the majority of expression differences are due to the gonad (Parisi et al. 2004), are in practice comparing expression related to sperm in males with a range of cell functions in females. Male-biased genes identified from whole-organism or bulk gonad preparations in many species show rapid rates of protein evolution (Ranz et al. 2003; Ellegren and Parsch 2007; Parsch and Ellegren 2013), and exhibit high rates of turnover (Zhang et al. 2007; Harrison et al. 2015; Papa et al. 2017; Whittle and Extavour 2019; Khodursky et al. 2020), both of which might be expected from sexual selection predictions. Indeed, in our bulk analysis, male-biased genes in the gonad show elevated rates of evolution (Fig. 3D). However, it is important to note that genes with restricted expression (e.g., restricted to sperm) often experience fewer adaptive constraints (Yannai et al. 2005) than those with broader expression. Indeed, when comparing genes that are sex-biased within gonad cell types, rather than those that differ as a result of cell-type abundance, male-biased genes in the gonad evolve at the same rate as unbiased genes (Fig. 3D).

Overall, our results suggest that sex differences in cell type abundance scale with visible sexual dimorphism, suggesting that bulk RNA-seq approaches may be more problematic in more dimorphic species. Single-cell transcriptomics, such as that employed here, offers a promising way to correct for any potential false inferences of differential gene expression and patterns of sequence divergence that are associated with standard bulk-level sequencing of heterogeneous tissue samples. However, although scRNA-sequencing approaches are increasingly employed for disentangling the role of regulatory changes in the evolution of intra- and inter-specific phenotypic variation (Fuess and Bolnick, 2021; Murat et al. 2021), the associated costs are still substantially higher compared to standard bulk RNA-sequencing and many technical challenges remain for non-model organisms (Alfieri et al. 2022). Cell isolation protocols are non-trivial as cell physiology may differ across tissues and species, and often require organism-specific technical knowledge to avoid cell death or bias in expression profiles (Reichard and Asosingh, 2019; Wang et al. 2021). Incomplete or poorly annotated genomes can additionally limit the identification of cell identities and gene expression patterns (Healey et al. 2022), although several *k*-mer based methods have been developed to aid with *de novo* transcriptome assembly and identification of cell populations in cases where a reference genome is lacking (Nip et al. 2020; Sun et al. 2021).

In the absence of scRNA-sequencing data or information regarding the cellular composition of tissues, we suggest that adopting more stringent fold change thresholds for calling differentially expressed genes has the potential to substantially reduce the false-positive rates associated with bulk-level sequencing of heterogeneous tissues. Although a few studies have accounted for increasing fold change thresholds when studying the evolution of differential gene expression (Mariani 2003; Perry et al. 2014; Harrison et al. 2015; Darolti et al. 2017; Ma et al. 2018; Ma et al. 2020; Djordjevic et al. 2022), others rely on a log_2_ fold change of 1 or lower, or on statistical significance alone. However, our results from bulk-level analyses show that this threshold results in the inclusion of many sex-biased genes that are due to differences in cell type abundance between males and females (Fig. S4). Although this attempt to reduce false-positive effects may also inadvertently remove some true-positive sex-biased genes, strong patterns of transcriptional dimorphism will remain unaffected. This approach would be particularly recommended for studies in which estimating additional tissue scaling parameters is limited due to the small size of the organism or difficulty in precise tissue dissection. Additionally, deconvolution methods may also prove useful (Monaco et al. 2019; Aguirre-Gamboa et al. 2020).

Taken together, our findings offer an important insight into the effect of allometry and cellular heterogeneity on inferred patterns of sex-biased gene expression and demonstrate the power of single cell RNA-sequencing in differentiating sex-biased genes that stem from regulatory rewiring between males and females from those that are due to differences in cell type abundance resulting from sex-specific developmental trajectories.

## Acknowledgments

We thank members of the Mank lab for helpful and constructive discussions on the project. This work was funded by grants from the European Research Council (grant number 680951), NSERC and CFI, as well as a Canada 150 Research Chair to J.E.M.

## Author contributions

I.D. and J.E.M. conceived the study, performed the analysis and wrote the manuscript.

## Conflict of interest

The authors declare no conflict of interest.

## Data accessibility

scRNA-sequencing reads and scripts used of data analysis will be made available upon publication.

## Notes

### Competing Interest Statement

The authors have declared no competing interest.

