## Supplementary Information for "Sex-biased gene expression at single-cell resolution: Cause and consequence of sexual dimorphism"

**Fig. S1. UMAP plot showing cell clustering of scRNA-seq datasets and identified cell populations in skin (A), heart (B), liver (C) and gonad (D).**

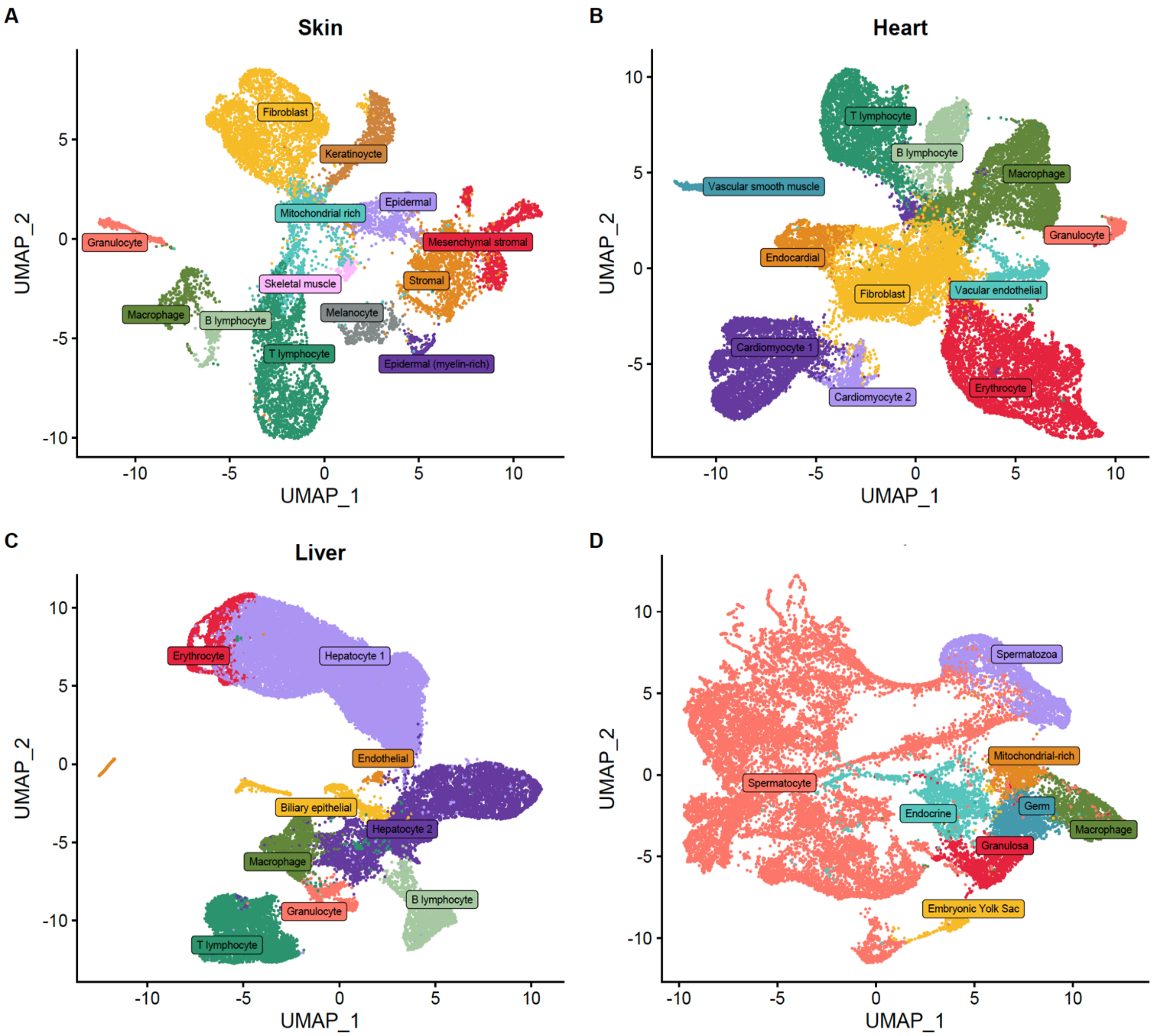

**Fig. S2. Marker gene expression across identified cell types of each tissue.**

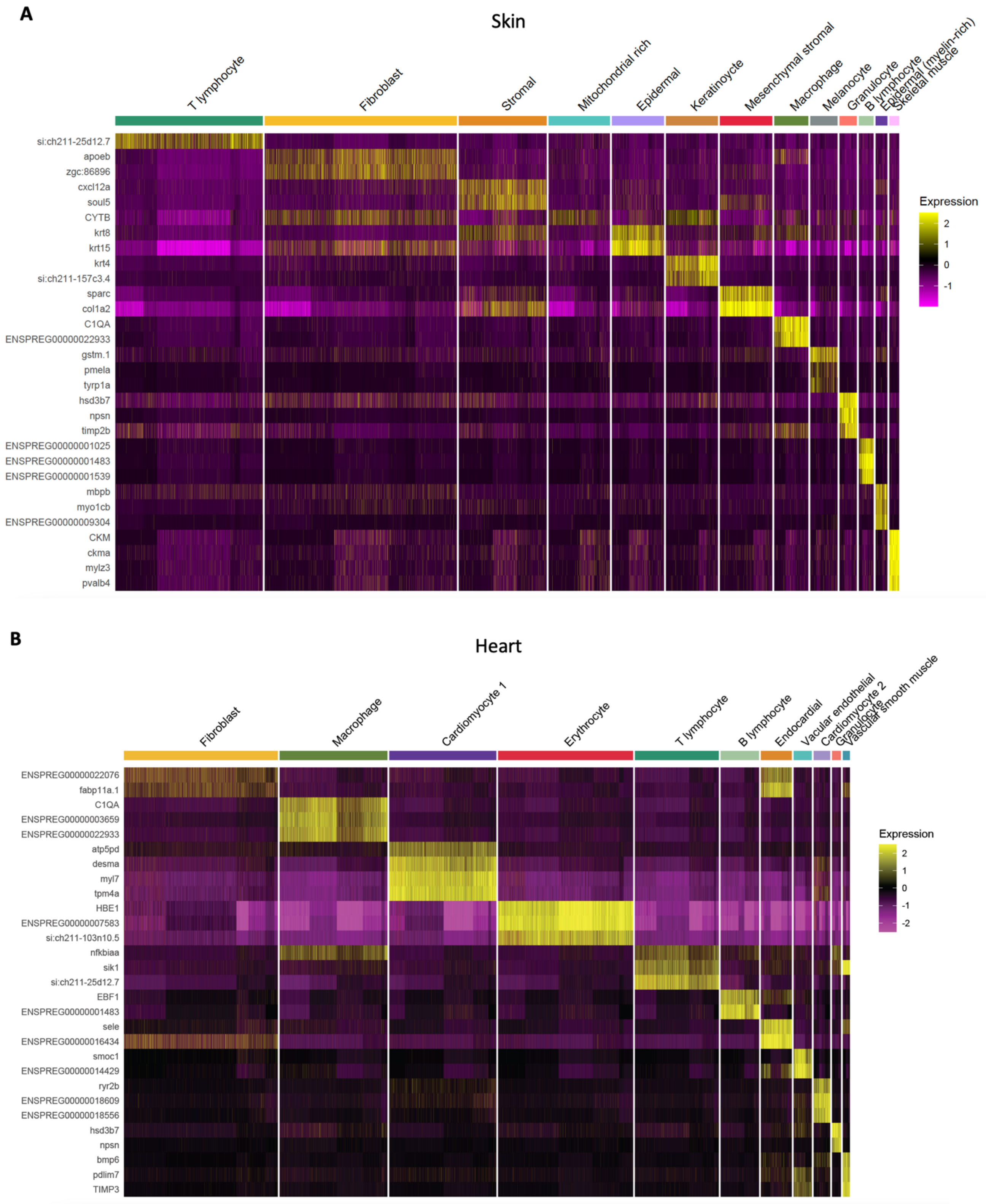

C

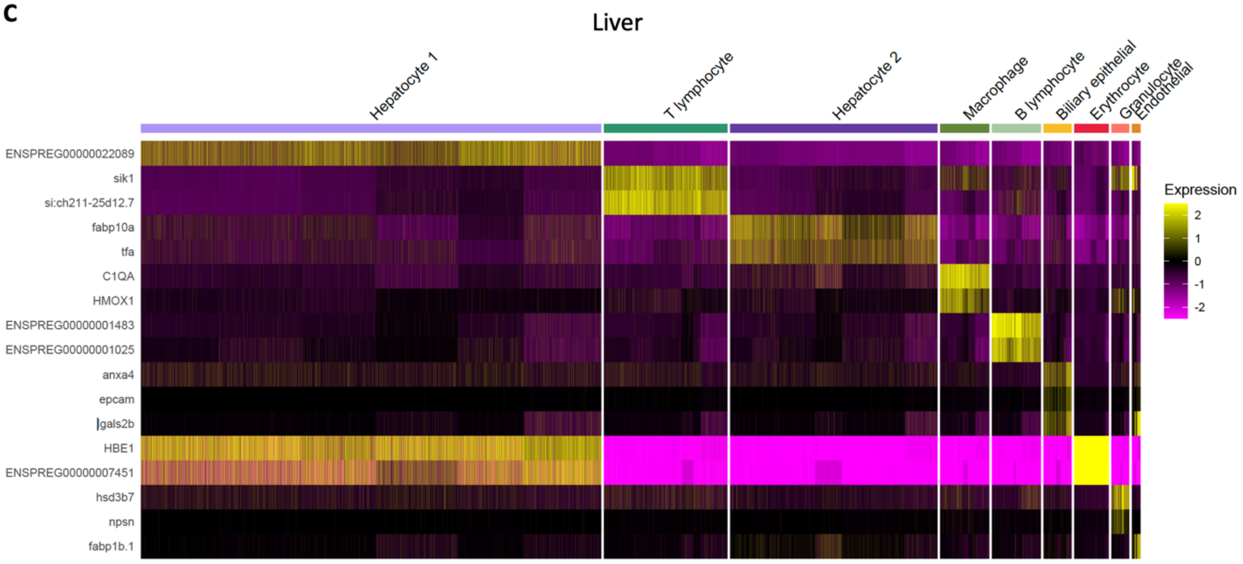
Liver
Hepatocyte 1
T lymphocyte
Hepatocyte 2
Macrophage
B lymphocyte
Biliary epithelial
Erythrocyte
Granulocyte
Endothelial
ENSPREG00000022089
sik1
si:ch211-25d12.7
fabp10a
tfa
C1QA
HMOX1
ENSPREG00000001483
ENSPREG00000001025
anxa4
epcam
lgals2b
HBE1
ENSPREG00000007451
hsd3b7
npsn
fabp1b.1
Expression
2
1
0
-1
-2

D

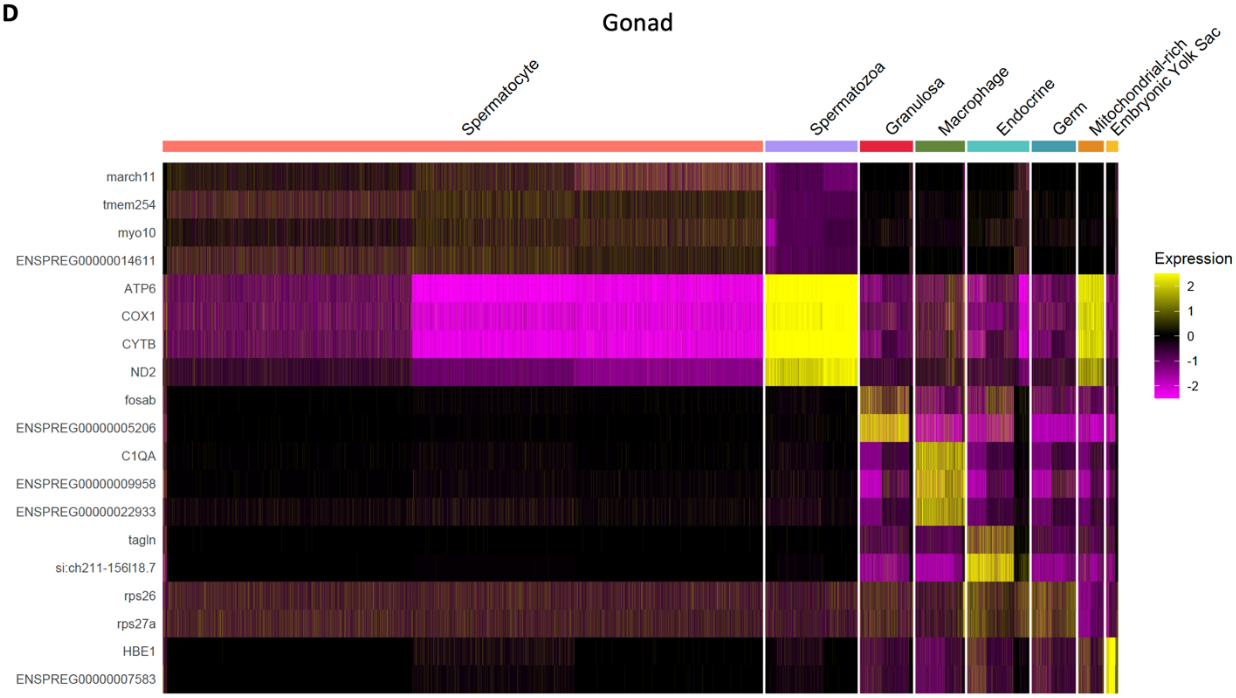
Gonad
Spermatocyte
Spermatozoa
Granulosa
Macrophage
Endocrine
Germ
Mitochondrial-rich
Embryonic Yolk Sac
march11
tmem254
myo10
ENSPREG00000014611
ATP6
COX1
CYTB
ND2
fosab
ENSPREG00000005206
C1QA
ENSPREG00000009958
ENSPREG00000022933
tagln
si:ch211-156l18.7
rps26
rps27a
HBE1
ENSPREG00000007583
Expression
2
1
0
-1
-2

**Fig. S3. Average male and female proportion of each identified cell type within each tissue.**

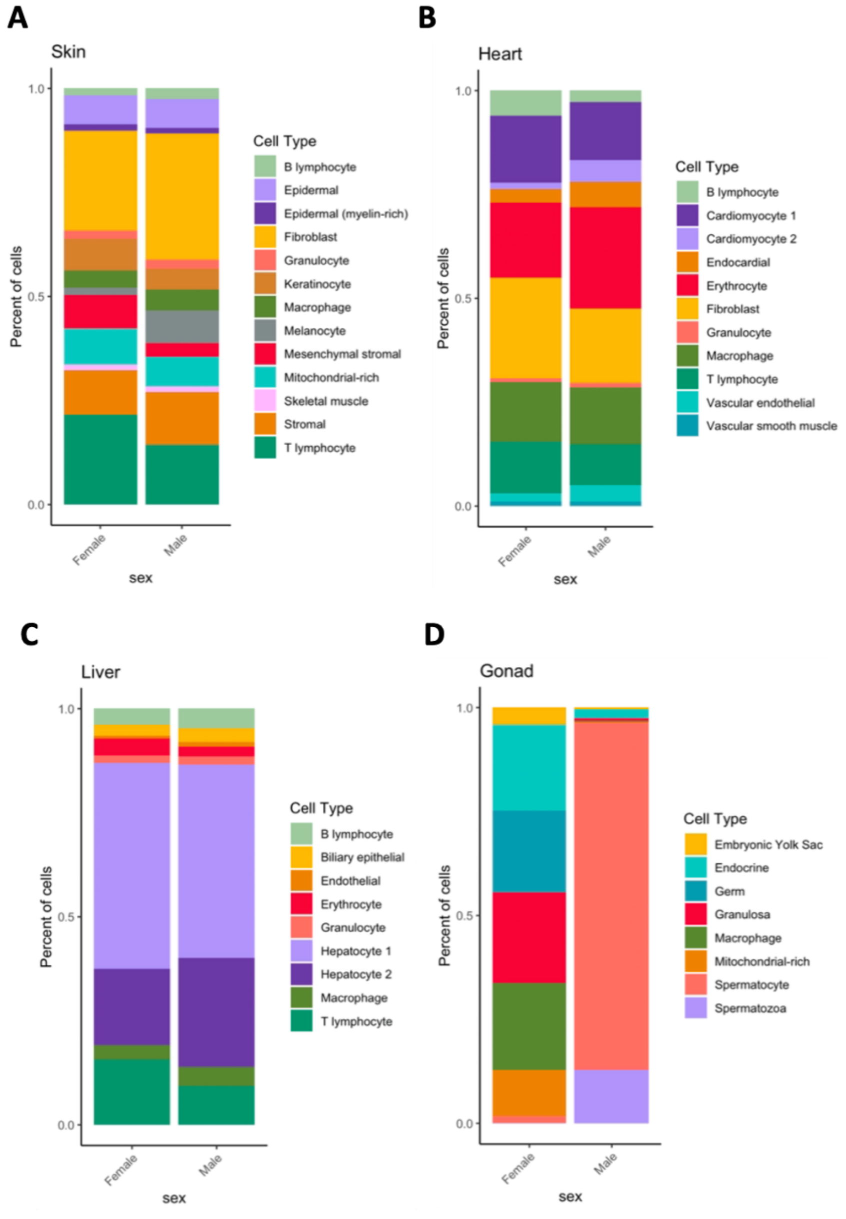

**Fig. S4. Magnitude of $log_2$ fold change in expression for female-biased (orange) and male-biased (green) genes.** Significance values are based on comparisons between genes with the same sex-bias direction, calculated using paired Wilcoxon's signed-rank tests (*** $p < 0.001$).

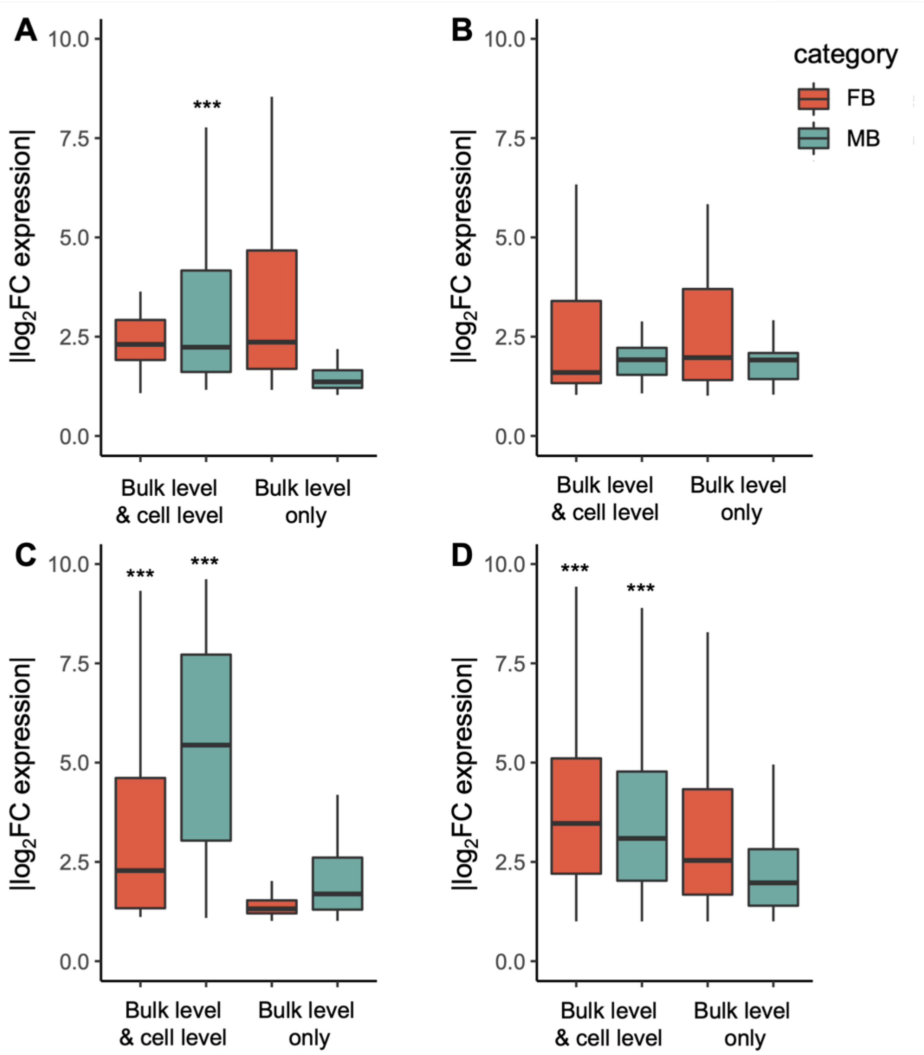

**Fig. S5. Expression (log₂ CPM) for each cell type within each tissue for genes identified as male-biased (green) and female-biased (orange) at the bulk level only.** Significance values are calculated using paired Wilcoxon's signed-rank tests (*** $p < 0.001$).

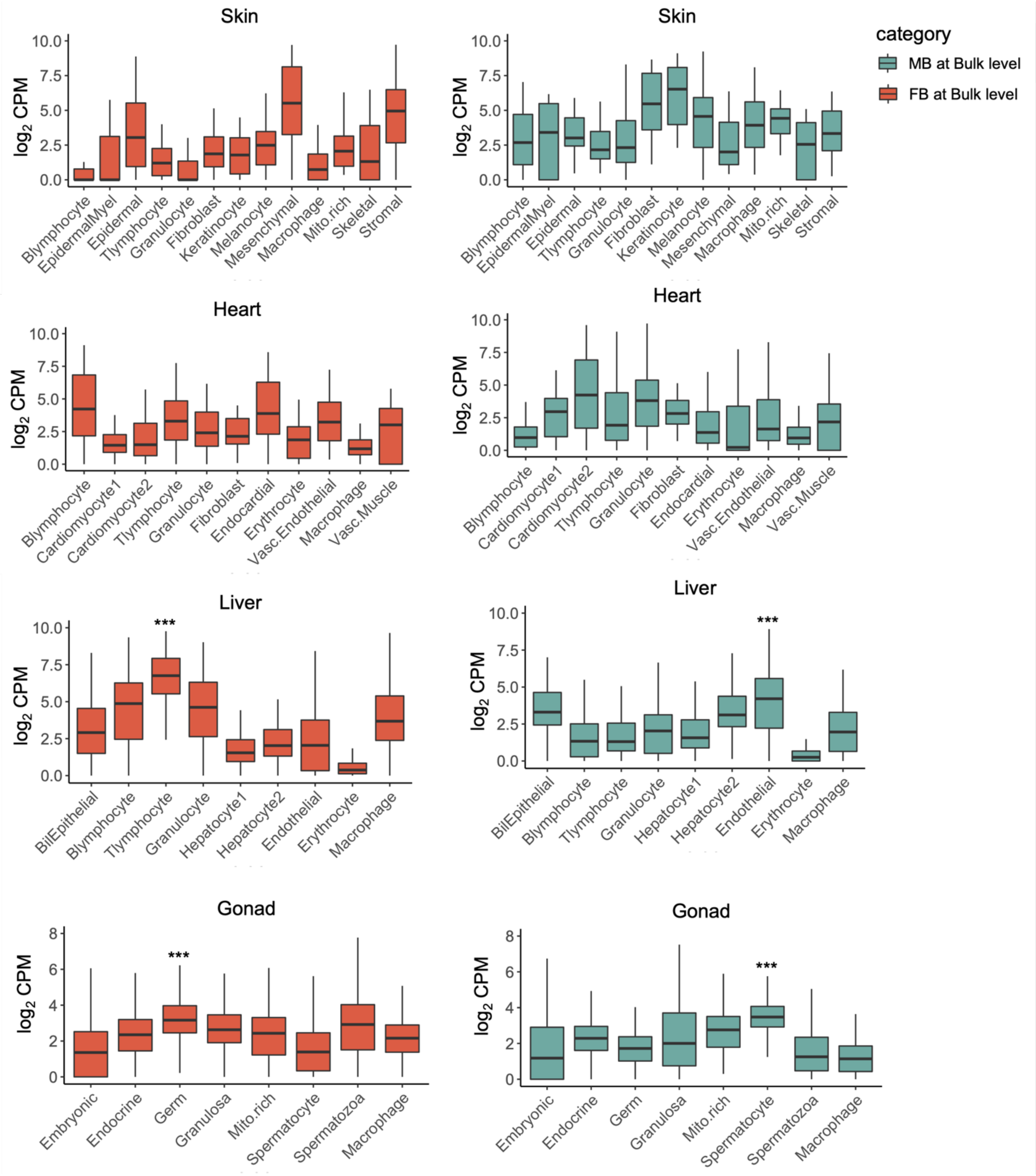

**Table S1. Differential cell type abundance between males and females.** Shown are male and female proportions for each identified cell type within each tissue and the $\log_2$ fold changes (female:male) in proportions. Significance based on two-proportions z-tests.

| Tissue | Cell Type | Female prop. | Male prop. | $\log_2$FC (F:M) | *p* value |
|---|---|---|---|---|---|
| **Skin** | Mesenchymal stromal | 0.082 | 0.033 | 1.286 | 0.001 |
| | T lymphocyte | 0.215 | 0.143 | 0.587 | 0.001 |
| | Keratinocyte | 0.076 | 0.051 | 0.584 | 0.001 |
| | Epidermal (myelin-rich) | 0.016 | 0.012 | 0.365 | 0.161 |
| | Mitochondrial-rich | 0.088 | 0.071 | 0.307 | 0.002 |
| | Epidermal | 0.070 | 0.070 | -0.014 | 0.920 |
| | Skeletal muscle | 0.013 | 0.013 | -0.018 | 1 |
| | Granulocyte | 0.021 | 0.022 | -0.064 | 0.790 |
| | Stromal | 0.107 | 0.128 | -0.254 | 0.001 |
| | Macrophage | 0.040 | 0.049 | -0.302 | 0.023 |
| | Fibroblast | 0.238 | 0.303 | -0.346 | 0.001 |
| | B lymphocyte | 0.017 | 0.025 | -0.547 | 0.005 |
| | Melanocyte | 0.017 | 0.079 | -2.178 | 0.001 |
| **Heart** | B lymphocyte | 0.060 | 0.027 | 1.120 | 0.001 |
| | Fibroblast | 0.243 | 0.179 | 0.446 | 0.001 |
| | T lymphocyte | 0.124 | 0.099 | 0.321 | 0.000 |
| | Cardiomyocyte 1 | 0.162 | 0.140 | 0.211 | 0.000 |
| | Macrophage | 0.144 | 0.136 | 0.076 | 0.129 |
| | Vascular smooth muscle | 0.010 | 0.010 | -0.015 | 0.994 |
| | Granulocyte | 0.009 | 0.011 | -0.277 | 0.163 |
| | Erythrocyte | 0.179 | 0.245 | -0.451 | 0.001 |
| | Endocardial | 0.033 | 0.062 | -0.915 | 0.001 |
| | Vascular endothelial | 0.020 | 0.039 | -0.947 | 0.001 |
| | Cardiomyocyte 2 | 0.016 | 0.051 | -1.676 | 0.001 |
| **Liver** | Erythrocyte | 0.043 | 0.024 | 0.830 | 0.001 |
| | T lymphocyte | 0.157 | 0.094 | 0.742 | 0.001 |
| | Hepatocyte 1 | 0.496 | 0.464 | 0.096 | 0.000 |
| | Granulocyte | 0.017 | 0.019 | -0.163 | 0.164 |
| | Biliary epithelial | 0.027 | 0.033 | -0.270 | 0.002 |
| | B lymphocyte | 0.038 | 0.046 | -0.277 | 0.000 |
| | Macrophage | 0.034 | 0.046 | -0.435 | 0.000 |
| | Hepatocyte 2 | 0.182 | 0.261 | -0.518 | 0.001 |
| | Endothelial | 0.005 | 0.012 | -1.254 | 0.001 |
| **Gonad** | Germ | 0.196 | 0.000 | 30.871 | 0.001 |
| | Mitochondrial-rich | 0.111 | 0.001 | 7.118 | 0.001 |
| | Macrophage | 0.210 | 0.003 | 6.017 | 0.001 |
| | Granulosa | 0.219 | 0.005 | 5.352 | 0.001 |
| | Embryonic Yolk Sac | 0.040 | 0.004 | 3.271 | 0.001 |
| | Endocrine | 0.207 | 0.022 | 3.250 | 0.001 |
| | Spermatocyte | 0.017 | 0.836 | -5.660 | 0.001 |
| | Spermatozoa | 0.001 | 0.129 | -7.760 | 0.001 |
